# EGFR-Driven Phenotypes Dictate Differential Therapeutic Response to Radiotherapy and Temozolomide in Glioblastoma

**DOI:** 10.1101/2025.11.22.689881

**Authors:** María Castelló-Pons, Miguel Perales-Patón, Matteo Italia, Irene Gómez-Soria, Alexis Casero, Juan Belmonte-Beitia, Víctor M. Pérez-García, Pilar Sánchez-Gómez

## Abstract

Despite the established Stupp regimen, glioblastoma (GBM) remains a highly lethal cancer with a 5-year survival rate below 10%. The epidermal growth factor receptor (EGFR) is frequently amplified or mutated in GBM, and we have previously shown that different EGFR statuses correlate with distinct tumor phenotypes and responses to temozolomide (TMZ). In this study, we investigated the differential response of two mouse GBM models, one overexpressing EGFRwt (EGFRwt/amp) and the other with overexpressing the EGFRvIII variant, to radiotherapy (RT) alone and in combination with TMZ. While both tumor models were sensitive to RT in vitro, in vivo experiments showed no significant survival benefit from RT for mice carrying EGFRwt/amp GBM, regardless of the RT schedule. Moreover, for these tumors, different combinations of RT with TMZ were not significantly better than spaced TMZ treatment alone. In contrast, EGFRvIII tumors responded well to RT alone, and spacing out the RT doses offered no additional benefit. Although a Stupp-like protocol provided a small benefit compared to RT alone in this model, the combinatorial treatment also induced the expression of several resistance markers, such as MGMT or NF-κb phosphorylation. Our retrospective analysis of patient data supports these findings, suggesting that RT alone may not improve survival for patients with EGFRwt/amp GBM, whereas for GBM with EGFRvIII mutations, adding RT or TMZ and RT, does provide a clear survival benefit to surgery or to surgery and RT, respectively. These results suggest that EGFR status could serve as a crucial biomarker to predict tumor response and guide personalized treatment decisions, particularly in cases where minimizing toxicity is a priority.

## Introduction

There has been very little progress in treating glioblastoma (GBM), a highly aggressive brain tumor, over the past several decades. The current standard of care, known as the Stupp regimen, was established in 2005 [1]. It involves maximal surgical resection followed by radiotherapy (RT; 60 Gy total, divided in 30 sessions) combined with concomitant temozolomide (TMZ), followed by six cycles of adjuvant TMZ. Despite this protocol, the population-based 5-year survival rate remains below 10% [2].

Multiple factors have been proposed as mediators of radioresistance in GBM, including DNA repair mechanisms, mesenchymal traits, and hypoxia. However, none of these have made it into routine clinical use. For chemotherapy, the only reliable biomarker for predicting a response to TMZ is the methylation status of the O6-methylguanine-DNA-methyltransferase (MGMT) promoter [3]. This enzyme reverses the cytotoxic effect of TMZ. Patients with an unmethylated MGMT promoter typically have a poor response to TMZ and are often treated with RT alone or enrolled in clinical trials.

The epidermal growth factor receptor (EGFR) gene is amplified in 50-60% of GBMs. Approximately half of these tumors also harbor the EGFRvIII variant, a deletion that leads to constitutive tyrosine kinase activation. In addition, missense mutations in the extracellular domain of EGFR are common. All these alterations enhance the tumor’s oncogenic potential and increase cell proliferation [4]. We formerly demonstrated that EGFR status also affects the vascular phenotype of these tumors. While EGFR wild type/amplified (EGFRwt/amp) tumors exhibit a highly angiogenic profile and blood-brain barrier (BBB) disruption, tumors with EGFRvIII or other point mutations have a more robust vasculature that fuel a rapid proliferative growth [5, 6]. These phenotypic differences influence both tumor progression and therapeutic response. For example, our previous work showed that an EGFRvIII GBM model did not respond to TMZ, whereas spaced doses of the drug slowed down EGFRwt/amp tumors and reduced resistance markers [7].

In this study, we analyzed the differential response of these two orthotopic mouse GBM models to RT alone and to the combination of TMZ and irradiation. We also used in silico models to optimize the treatment protocols. We found that, although both EGFRwt-amp and EGFRvIII cells respond to RT in vitro, EGFRwt-amp tumors did not show a significant response to irradiation in vivo. For EGFRvIII tumors, there was no benefit on spacing out the RT doses. Regarding combinatorial treatments, for mice carrying EGFRwt/amp cells, the combination of RT with TMZ was not significantly better than spaced TMZ alone. In contrast, for EGFRvIII tumors, a Stupp-like protocol provided a small benefit compared to RT alone, even though the combinatorial treatment induced the expression of several resistance markers, such as MGMT or NF-κB phosphorylation [8]. Our retrospective analysis of patient data supports these findings, suggesting that RT alone may not improve survival for patients with EGFRwt/amp GBM. By contrast, for GBM with EGFR mutations, adding RT or TMZ and RT, does provide a clear survival benefit to surgery or to surgery and RT, respectively. These results suggest that EGFR status could be a key predictor of tumor response to RT and TMZ, helping to identify the most effective treatment approach, particularly for elderly patients where toxicity is a major concern.

## Materials and Methods

### In vitro assays

#### GBM cell lines

Two mouse-derived GB cell lines were used in this study: SVZ-wt and SVZ-vIII. Both cell lines were generated and characterized in our laboratory previously. They were derived from p16/p19 knockout subventricular zone (SVZ) progenitors following the overexpression of either wild-type EGFR (EGFRwt) or the truncated isoform EGFRvIII, and both cell lines constitutively express the luciferase reporter gene [9]. The cells were cultured in Complete Medium: DMEM/F-12 medium (Corning) supplemented with B27 (1:50; Gibco), penicillin-streptomycin (1%; Lonza), ciprofloxacin (2 mg/ml), heparin (0.2%; Sigma-Aldrich), and the growth factors EGF (20 ng/ml) and FGF (10 ng/ml; Peprotech). The cultures were maintained at 37 °C with 5% CO₂ and were routinely passaged using Accumax (20 µl/ml, Millipore).

#### DNA damage analysis after irradiation

SVZ-EGFRwt and SVZ-EGFRvIII cells (3 × 10⁴) were seeded on coverslips coated with Matrigel® Matrix (Corning) in 24-well plates and irradiated 24 hours later with 1, 2, or 5 Gy using a MultiRAD 225 X-ray system (Faxitron). Two hours after irradiation, the cells were fixed with 4% paraformaldehyde (PFA; 15 minutes, room temperature), incubated with anti-γH2AX antibody at a dilution 1:100 overnight at 4 °C (Cell Signaling Technology). This was followed by incubation with an Alexa Fluor 647-conjugated secondary antibody at a dilution of 1:200 for 1 hour at room temperature (Thermo Fisher Scientific). Nuclei were counterstained with DAPI (5 µg/ml; Sigma-Aldrich), and the coverslips were mounted on glass slides using Fluoromount aqueous mounting medium (SouthernBiotech). Images were acquired using a Hamamatsu Orca-Spark camera coupled to a Leica fluorescence microscope.

#### Cell viability assays

SVZ-EGFRwt and SVZ-EGFRvIII cells (1 × 10⁴ per well) were seeded in triplicate in 96-well plates and irradiated 24 hours later with 1, 2, or 5 Gy. Cell viability was assessed by two complementary methods: (i) bioluminescence measurement 24 hours post-irradiation after a 15min incubation with D-luciferin (15 µg/ml, 1:100; PerkinElmer) using a Spark TECAN microplate reader (exposure time: 1000 ms); and (ii) an MTT assay (3 mg/ml, 1:2; Thermo Fisher Scientific) performed 48 hours post-irradiation after 1-hour incubation, with absorbance measured at 570 nm. Representative images of irradiated cells were acquired using a Hamamatsu camera coupled to a Leica microscope.

### In vivo assays

#### Orthotopic mouse glioma models

All animal experiments were approved by the Research Ethics and Animal Welfare Committee (PROEX 306.7/22) at Instituto de Salud Carlos III (Madrid, Spain) and were conducted in accordance with European and national regulations. Orthotopic gliomas were generated by stereotaxic injection of 2.5 × 10⁵ cells in 2 μl of complete medium into the brains of nude mice at the following coordinates relative to Bregma: A–P −0.5 mm; M–L +2 mm; D–V −3 mm (Leica stereotaxic device). Three days after implantation, the mice underwent treatment according to the different experimental schedules, which included RT (3 Gy/dose, using a MultiRAD 225 X-ray system (Faxitron) with a 0.5 mm Cu filter), a combination of RT and TMZ (10 mg/kg/dose; MedChemExpress) administered intraperitoneally (IP) after dilution in PBS containing 1% bovine serum albumin (BSA), or TMZ alone. Control animals received the corresponding solvent. Tumor growth was monitored using bioluminescence imaging (IVIS, PerkinElmer) following the IP of D-luciferin (75 mg/kg; Thermo Fisher Scientific). The animals were euthanized when they showed signs of tumor growth, and brains were collected and freshly frozen for cellular and molecular analyses.

#### Quantitative reverse-transcriptase PCR (qRT-PCR)

Total RNA was extracted from frozen glioma tissues using TRIzol Reagent® (Applied Biosystems) following the manufacturer’s instructions. The concentration and purity of the RNA were determined using a NanoDrop One spectrophotometer (Thermo Scientific). Complementary DNA (cDNA) was synthesized from 1 μg of RNA using a PrimeScript RT Reagent Kit (Takara). Quantitative real-time PCR (qRT-PCR) was performed on a LightCycler 480 system (Roche Diagnostics) using TB Green Premix Ex Taq™ II (Takara) and specific primers (Sigma-Aldrich) for the genes of interest (Supplementary Table S1). Relative gene expression was calculated using the 2^(−ΔΔ) Ct method, with β-actin serving as the reference gene.

#### Western Blot (WB) analysis

Protein extracts were obtained from glioma tissue samples by resuspending the samples in lysis buffer (50 mM de Tris (pH 7.6), 400 mM de NaCl, 1% de SDS, 1mM EDTA, 1mM EGTA) and incubating for 15 minutes at 100 °C. The lysates were then centrifuged at 13,000 g for 10 minutes at RT, and protein concentration was determined using a BCA Protein Assay Kit (Thermo Fisher Scientific). Equal amounts of protein (30 μg per sample) were separated using 12% SDS-PAGE gels in running buffer (25 mM Tris base, 192 mM glycine, 0.1% SDS) and transferred to nitrocellulose membranes (Hybond-ECL, Amersham Biosciences, GE Healthcare) using the Bio-Rad Mini Trans-Blot system (80 V, 2 hours, 4 °C). The membranes were blocked for 1 hour at RT in 5% non-fat dry milk prepared in TBS-T (10 mM Tris-HCl (pH 7.5), 100 mM NaCl, 0.1% Tween-20).

The membranes were subsequently incubated overnight at 4 °C with the following primary antibodies diluted in blocking buffer: anti-MGMT (1:1000, Cell Signaling Technology), phospho–NF-κB p65 (Ser536) (1:1000, Cell Signaling Technology), and anti-Rho (1:800, Santa Cruz Biotechnology). After washing, membranes were incubated with HRP-conjugated secondary antibodies (anti-rabbit IgG, 1:2500, anti-mouse IgG, 1:2500; Cell Signaling Technology). Proteins were visualized using enhanced chemiluminescence (ECL, Pierce) with the ImageQuant 800 system (Amersham Biosciences), and band intensity was quantified using ImageJ software (Cytiva).

#### Statistical Analysis

Data were represented and analyzed statistically using GraphPad Prism 8.0 software (GraphPad Software, Inc.). The results are presented as mean comparisons, and the Kolmogorov–Smirnov test was used to assess variable normality. For comparisons involving more than two groups, one-way ANOVA was applied; in cases of non-normality, the non-parametric Kruskal–Wallis test was used instead. Survival curves were generated using the Kaplan–Meier method, and group differences were evaluated using the Mantel–Cox log-rank test. Statistical significance was set at p < 0.05, with probabilities represented as follows: * p < 0.05, ** p < 0.01 and *** p < 0.001.

### Mathematical model

We employed a mathematical model of GBM growth under chemoradiotherapy that has been previously described, calibrated, and validated (Perales-Patón et al., under review). This model was adopted for two main reasons. First, it is based on the same mouse model investigated in the present study. Second, it enables the generation of virtual mouse cohorts by estimating individual-specific parameters—derived from experimental data or the literature—and simulating tumor response under alternative therapeutic regimens, as detailed in the original work.

As an initial step, the model was used to systematically explore treatment schedules involving fixed RT and TMZ doses administered at equidistant intervals. Two timing variables were varied across simulations: (i) the delay between the initiation of the two treatments (ranging from 0 to 14 days), and (ii) the spacing between consecutive treatment administrations (ranging from 1 to 14 days). Beyond these constrained schemes, additional protocols were explored, including metronomic schedules and a mouse-adapted, Stupp-like regimen.

The metronomic schedules were designed to evaluate smaller TMZ doses delivered as single injections (3.33 mg/kg in vivo per administration) while maintaining the same cumulative drug exposure. In silico, equidistant inter-dose intervals were progressively varied to identify the regimen yielding the greatest survival benefit, either as monotherapy or in combination with RT. These simulations indicated that RT delivered concomitantly with the first three TMZ doses, followed by TMZ administered at regular intervals, provided the greatest improvement in overall survival. Median OS increased as the inter-dose spacing widened, reaching a maximum at 11 days, beyond which efficacy declined. To ensure laboratory feasibility, we selected a 7-day spacing for in vivo implementation. This corresponded to weekly TMZ injections for nine weeks, with 3 Gy RT delivered once weekly during the first three weeks.

The Stupp-like protocol was adapted to reproduce the clinical standard of care for human patients, as previously described (Perales-Patón et al., under review). It consisted of daily 3 Gy RT with concomitant TMZ for three consecutive days (one injection per day, corresponding to 3.33 mg/kg in vivo), followed by a two-day break and three additional adjuvant TMZ doses administered every three days (two injections per dose, corresponding to 6.66 mg/kg in vivo). Finally, TMZ monotherapy administered at 14-day intervals (TMZ ×14), previously associated with favorable in vivo outcomes [7], was also simulated.

### Retrospective patient’s data (TOG)

The data used in this study were derived from TOG (Therapy Optimization in Glioblastoma), a collaborative project that collects and curates clinical and molecular information from GBM patients across several Spanish hospitals. Specifically, we retrospectively analyzed 336 newly diagnosed GBM patients treated between 2010 and 2020 across six centers: 103 from University Hospital 12 de Octubre (Madrid), 70 from Virgen de la Salud Hospital (Toledo), 55 from Ciudad Real University General Hospital (Ciudad Real), 51 from Marqués de Valdecilla University Hospital (Santander), 46 from Carlos Haya Regional University Hospital (Málaga), and 11 from Manises Clinical Hospital (Valencia). Collected variables included patient age, sex, Karnofsky Performance Status (KPS) at treatment initiation, type of surgery—gross total resection (GTR), partial resection (PR), or biopsy (B)—, treatment modality (surgery only, RT only, TMZ only, or CRT following the Stupp protocol), molecular biomarkers (EGFR, IDH, MGMT, and Ki67), and overall survival (OS). The primary study endpoint was the OS, defined as the time from diagnosis to death. OS was analyzed using Kaplan-Meier survival curves, and survival differences between groups were assessed using previously described methodology (details are given in Suppl. Table 1).

Molecular characterization was performed following standard protocols. Ki67 expression, MGMT promoter methylation, IDH and EGFR mutation status were determined by immunohistochemistry (IHC). Patients with available EGFR data were stratified by treatment group (chemo and radiotherapy (CRT), RT only, or surgery only, see Suppl. Table T1). Baseline demographic, clinical, surgical, and molecular characteristics are summarized in Table 1.

**Table 1.** Clinical and molecular characteristics of patients (n = 136; selected cohort) stratified by treatment.

| Characteristics |  | Only surgery<br>(n=17) | Treated with RT (n=13) | Treated with CRT (n=106) |
| --- | --- | --- | --- | --- |
| Age, years | Median [interval] | 72 [42,86] | 70 [36,80] | 61 [24,83] |
|  | <70 (%) | 8 (47.1) | 6 (46.1) | 82 (77.4) |
|  | >=70 (%) | 9 (52.9) | 7 (53.9) | 24 (22.6) |
| Sex | Female (%) | 3 (18.8%) | 6 (46.1) | 46 (43.4) |
|  | Male (%) | 14 (81.2%) | 7 (53.9) | 60 (56.6) |
| KPS | Median [interval] | 70 [50,100] | 90 [50,100] | 80 [60, 100] |
|  | <70 (%) | 7 (41.1) | 2 (15.4) | 15 (14.1) |
|  | >=70 (%) | 10 (51.8) | 11 (84.6) | 91 (85.9) |
| Extent of resection | Biopsy | 10 (58.8) | 6 (46.2) | 28 (26.4) |
|  | Partial | 5 (29.4) | 4 (30.8) | 35 (33) |
|  | Gross total | 2 (11.8) | 3 (23) | 43 (45.6) |

| Characteristics | Only surgery<br>(n=17) | Treated with RT (n=13) | Treated with CRT (n=106) |
| --- | --- | --- | --- |
| **IDH mutation status** |  |  |  |
| Yes | 0 (0) | 1 (7.7) | 6 (5.7) |
| No | 17 (100) | 12 (92.3) | 100 (94.3) |
| ND | 0 (0) | 0 (0) | 0 (0) |
| **EGFR mutation status** |  |  |  |
| Yes | 7 (41.2) | 7 (53.9) | 57 (53.8) |
| No | 9 (52.9) | 6 (46.2) | 49 (46.2) |
| ND | 1 (5.9) | 0 (0) | 0 (0) |
| **MGMT promoter status** |  |  |  |
| Methylated | 4 (23.5) | 7 (53.9) | 37 (34.9) |
| Unmethylated | 4 (23.5) | 2 (15) | 25 (23.6) |
| ND | 9 (52.9) | 4 (30.1) | 44 (41.5) |

### Retrospective patient’s data

The Cancer Genome Atlas (TCGA) GBM cohort was accessed through the TCGA Data Portal (https://portal.gdc.cancer.gov/) to extract data on OS, EGFR mutation status, and treatment received. TCGA provides a large, publicly available, and extensively curated dataset from a multicenter patient population.

Patients were first stratified according to their EGFR status (wild type vs. mutant). Two initial groups were defined within each stratum: patients who had received RT (with or without TMZ) and patients who had received TMZ (with or without RT). The “set operations” tool available in the TCGA Data Portal was then applied to determine the intersection and difference between these groups, thereby defining two final subgroups: patients who received CRT (RT plus TMZ) and those treated with RT only. Kaplan–Meier survival curves were generated for each subgroup, and survival differences were assessed as described in the Statistical Analysis section.

### Results

#### In vitro response of EGFRwt and EGFRvIII cells to irradiation

In this study, we evaluated the in vitro response of two mouse GBM models to RT. These models were previously generated in our lab by overexpressing either EGFRwt or EGFRvIII in p16/p19 knockout subventricular zone (SVZ) progenitors. When injected intracranially into nude mice, both cell lines consistently generated gliomas, but the SVZ-EGFRwt tumors were less aggressive than the SVZ-EGFRvIII tumors. Moreover, tumors formed by SVZ-EGFRvIII cells were more proliferative than those from SVZ-EGFRwt cells, while the latter exhibited greater BBB disruption and a more hypoxic phenotype [5].

To test if EGFR status affects the sensitivity of GBM cells to RT, we treated in vitro both SVZ-EGFRwt and SVZ-EGFRvIII cells with increasing doses of irradiation (1, 2, and 5Gy) and assessed the formation of DNA-repair foci. We observed punctuated phospho-γH2AX staining in both cell lines 2 hours after 1 Gy and 2 Gy irradiation, with 5 Gy inducing a more diffuse and bright nuclear stain (Figure 1A-B). These observations were consistent with our cell viability studies. We found that 24 hours after irradiation, the cell number was reduced in both SVZ-EGFRwt (Fig. 1C, E) and SVZ-EGFRvIII (Fig. 1D, G) cells at all doses tested. The reduction in cell viability was even more pronounced 48 hours after exposure to 5 Gy. These results suggest that RT induces DNA damage in both cell lines, with higher doses leading to a significant reduction in cell number regardless of the presence of EGFR mutations.

**Figure 1.**
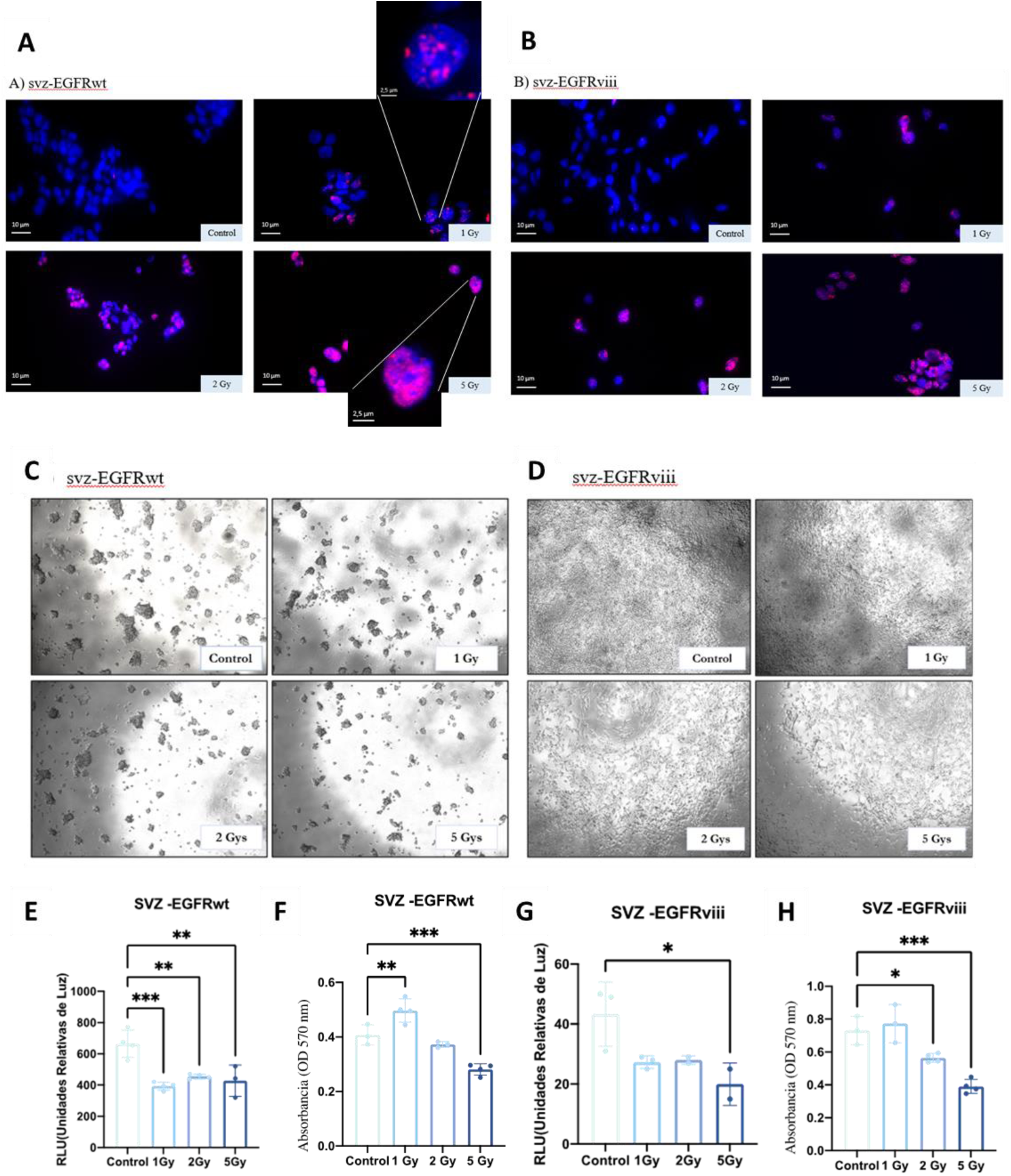
In vitro response of SVZ-EGFRwt and SVZ-EGFRvIII cells to irradiation. **(A-B)** Representative immunofluorescence images of the cells stained with anti-phospho γ-H2AX (DNA damage response foci) (magenta) 2h after RT treatment (0 (Control), 1, 2, and 5 Gy). DAPI (blue) was used to counterstain the nuclei. **(C-D).** Phase-contrast images of SVZ-EGFRwt **(C)** and SVZ-EGFRvIII **(D)** 24 h after irradiation. **(E, G)** Quantification of cell viability by bioluminescence 24 h after RT following D-luciferin incubation. **(F, H)** Quantification of cell viability by MTT assay performed 48 h after RT. *p < 0.05, **p < 0.01, *p < 0.001.

#### In vivo response of EGFRwt and EGFRvIII tumors to irradiation

Our previous findings showed that while both cell lines are sensitive to TMZ, SVZ-EGFRvIII tumors do not respond to this chemotherapy in vivo. In contrast, SVZ-EGFRwt tumor growth is inhibited by the drug when administered with sufficient spacing between doses [7]. We designed a similar experiment to evaluate the in vivo response of both models to different RT schedules. For that, we injected SVZ-EGFRwt and SVZ-EGFRvIII cells into the brains of immunodeficient mice. The animals received three doses of RT (3 Gy each) administered on consecutive days (X+1), or every 4 (X+4) or 7 (X+7) days (Fig. 2A). The survival of mice with SVZ-EGFRwt tumors was not affected by any of the RT schedules (Fig. 2B), but all treatments decreased the SVZ-EGFRvIII tumor burden (Fig. 2C). These results suggest that SVZ-EGFRwt cells are radioresistant within the tumor microenvironment, whereas the phenotype of SVZ-EGFRvIII tumors does not provide the same protection against irradiation. Notably, we did not observe significant changes in the expression of a panel of genes related to tumor phenotype (mesenchymal, proneural, vascular, persister) or radio-resistance in SVZ-EGFRvIII tumors treated with the different RT schedules (Supp. Fig. S1). This suggests that, unlike our previous results with TMZ [7], irradiation does not induce an apparent permanent phenotypic change in the presence of the EGFR deletion.

**Figure 2.**
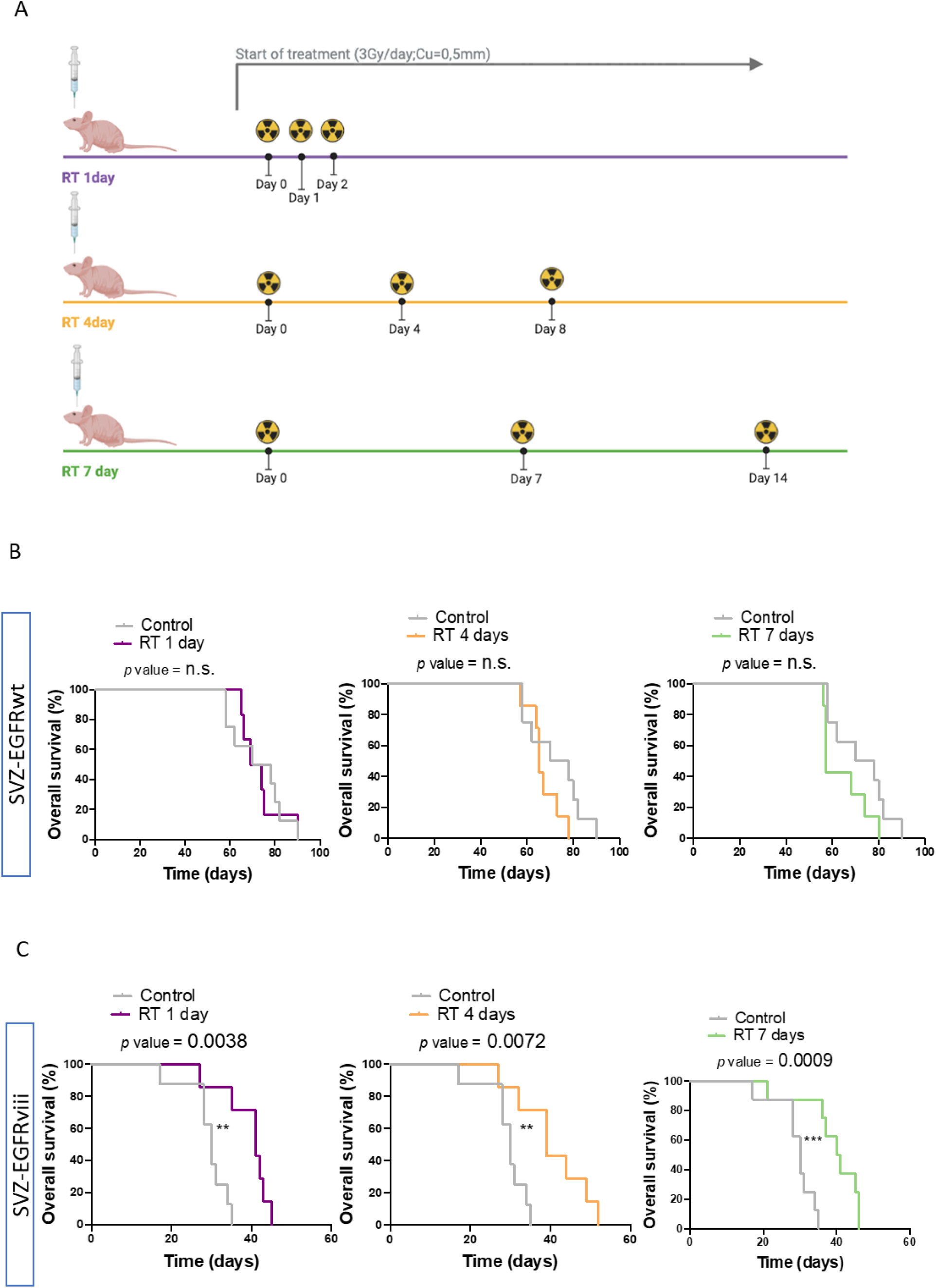
In vivo response of SVZ-EGFRwt and SVZ-EGFRvIII tumors to different RT schedules. **(A)** Schematic representation of the radiotherapy (RT) regimens, consisting of three fractions of 3 Gy delivered on consecutive days (X+1), or spaced every 4 days (X+4) or 7 days (X+7). **(B-C)** Kaplan-Meier overall survival curves of mice that were orthotopically injected with SVZ-EGFRwt **(B)** or SVZ-EGFRvIII **(C)** cells and subsequently treated with three RT doses (3 Gy, Cu=0.5 mm) with the different schedules. *p < 0.05, **p < 0.01, *p < 0.001.

#### In vivo response of EGFRwt and EGFRvIII tumors to combinatorial RT and TMZ treatment

To emulate the Stupp protocol used for GBM patients, we designed a combinatorial treatment regimen. It included three doses of RT (3 Gy each) on consecutive days, delivered concurrently with TMZ treatment, followed by three cycles of adjuvant TMZ (Fig. 3A). This Stupp-like treatment reduced tumor growth in both models (Fig. 3A-B). We observed an increased survival benefit for animals with SVZ-EGFRvIII tumors compared to those receiving RT alone. By contrast, the Stupp-like protocol provided a similar benefit to the one we previously observed in response to chemotherapy alone for the SVZ-EGFRwt model alone [7].

**Figure 3.**
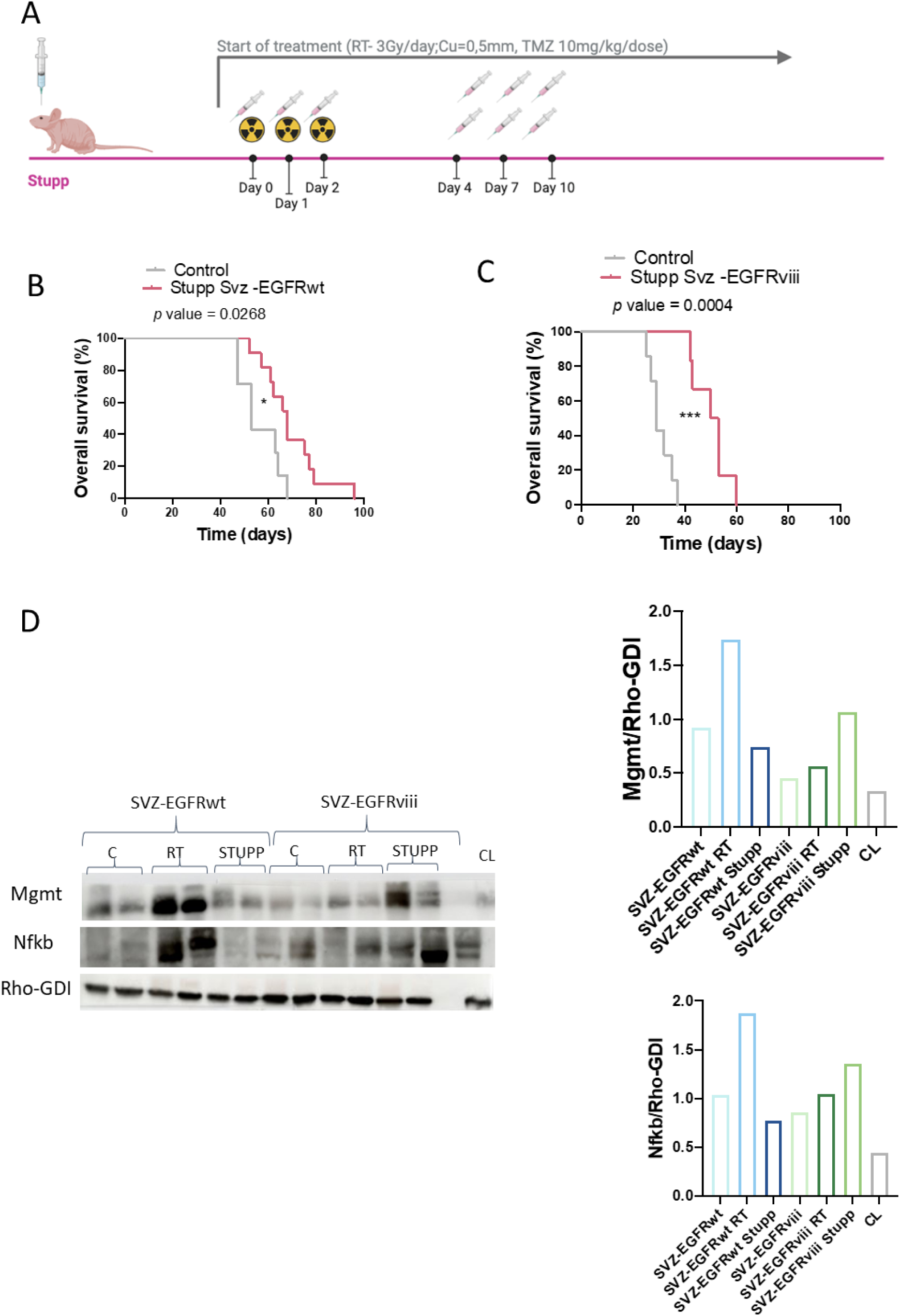
Stupp-like treatment scheme and analysis of SVZ-EGFRwt and SVZ-EGFRvIII tumors. **(A)** Schematic representation of Stupp-like protocol. **(B-C)** Kaplan–Meier overall survival curves of mice orthotopically injected with SVZ-EGFRwt **(B)** or SVZ-EGFRvIII **(C)** cells and subsequently treated with 9 intraperitoneal injections of TMZ (10 mg/kg per injection) and 3 RT doses (3 Gy, Cu = 0.5 mm) following the scheme. **(D)** Western blot analysis of MGMT (23 kDa) and phospho-NFkB (65kDa) expression in SVZ-EGFRwt and SVZ-EGFRvIII tumors treated with RT or the Stupp-like protocol along with their respective untreated tumors. Rho-GDI (23 kDa) was used as a loading control. Quantification is shown on the right. *p < 0.05, ***p < 0.001.

We also measured the expression of proteins potentially related to chemo- and/or radio-resistance, such as MGMT and phospho-NFκB. We observed that RT (X+1) induced a strong accumulation of both proteins in SVZ-EGFRwt tumors. For SVZ-EGFRvIII tumors, no clear induction was observed with RT alone, although a small increase was noted in tumors that received the Stupp-like protocol (Fig. 3D).

#### Using in silico models to optimize the combinatorial treatment of SVZ-EGFRwt tumors

In our previous work (Perales-Patón et al, under review), we reported *in silico* simulations of the Stupp-like, TMZ x+14, and CRT x+14 protocols (three daily sessions of 3 Gy RT combined with 30 mg/kg TMZ, each administered 14 days apart. Among these, the CRT x+14 regimen yielded the most favorable median OS outcome within the constrained search space of schedules in which RT and TMZ doses were administered at equidistant and with constant doses. In the present study, these regimens were included as reference benchmarks to enable comparison with additional schedules (Fig. 4A). The computational analysis was extended to incorporate the metronomic schedule, which had not previously been explored. This protocol produced *in silico* survival outcomes comparable to those of the more intensive Stupp-like protocol (Fig. 4B).

**Figure 4.**
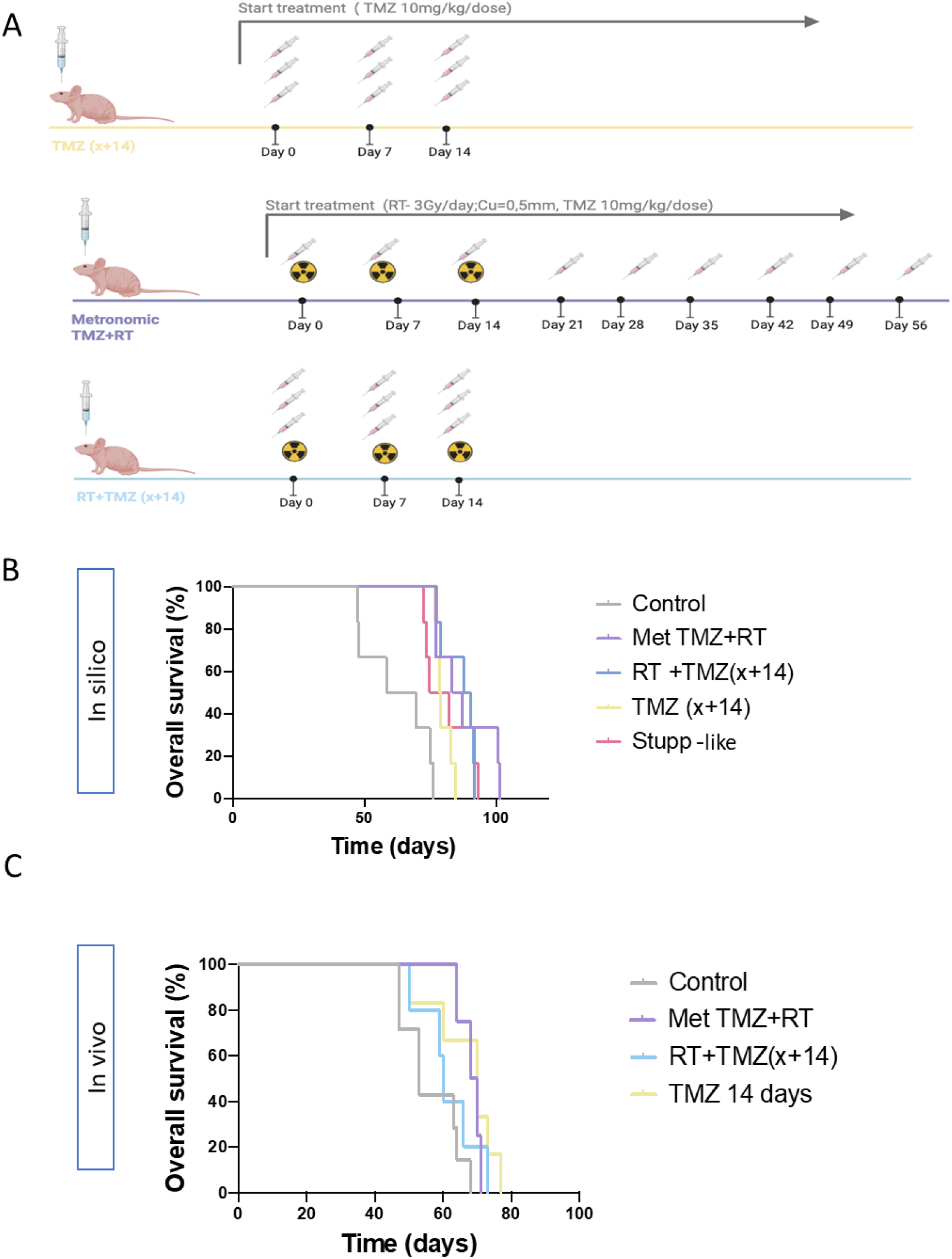
In silico and in vivo evaluation of treatment schedules for SVZ-EGFRwt tumors. **(A)** In silico simulations of the Stupp-like, RT+TMZ ×14, TMZ ×14, and metronomic regimens generated with the mathematical model. Survival curves are shown for all schedules. **(B)** In vivo Kaplan–Meier overall survival curves of mice orthotopically injected with SVZ-EGFRwt cells and treated with Stupp-like, RT+TMZ ×14, TMZ ×14, or the metronomic regimen.

Guided by these simulations, and aiming to improve OS outcomes beyond those of the Stupp-like protocol while using a less intensive regimen, four protocols were subsequently evaluated *in vivo*: the Stupp-like protocol—used as the control group in this experiment—the metronomic schedule, CRT x+14, and TMZ x+14. None of the alternative strategies achieved a statistically significant extension of survival compared with the Stupp-like protocol (median survival: metronomic, 68 days; CRT x+14, 60 days; TMZ x+14, 70 days; Stupp-like, 66 days) (Fig. 4C). Nevertheless, TMZ x+14 monotherapy yielded outcomes comparable to all CRT regimens, including the Stupp-like protocol (p = 0.789 for Stupp-like vs. TMZ x+14), and the addition of concomitant RT to TMZ x+14 (i.e., CRT x+14) did not provide any survival benefit (Fig. 4C).

#### Retrospective analysis of human data

Our aim was to evaluate, in the TOG cohort, whether chemotherapy alone could achieve OS rates comparable to the combination of TMZ with RT in EGFRwt patients, as suggested by in silico and in vivo models. Importantly, such an effect was not observed in EGFRmut tumors, where these models indicated that CRT outperformed both RT and TMZ monotherapy. However, because of the scarcity of patients treated with TMZ alone (N=5), direct CRT vs. TMZ comparisons could not be done. Instead, we compared RT monotherapy with CRT and surgery-only cases to evaluate the independent effect of RT and the added benefit of TMZ, while analyzing the prognostic role of EGFR.

When stratified by EGFR status, clear differences emerged. In EGFRwt tumors, RT alone was not effective, yielding outcomes similar to surgery (median OS 2.5 vs. 4.6 months, respectively; p = 0.64), whereas CRT significantly improved survival compared with both (median OS 10.5 months; p = 0.002 vs. RT) (Fig. 5), consistent with the results in mouse models. In EGFRmut tumors, RT alone improved survival relative to surgery (median OS 6.4 vs. 2.3 months; p = 0.03), and CRT further prolonged survival compared with RT alone (median OS 13.1 months; p = 0.016) (Fig. 5). These findings mirror our preclinical evidence and reinforce the idea that EGFR status modulates the efficacy of RT and CRT.

**Figure 5.**
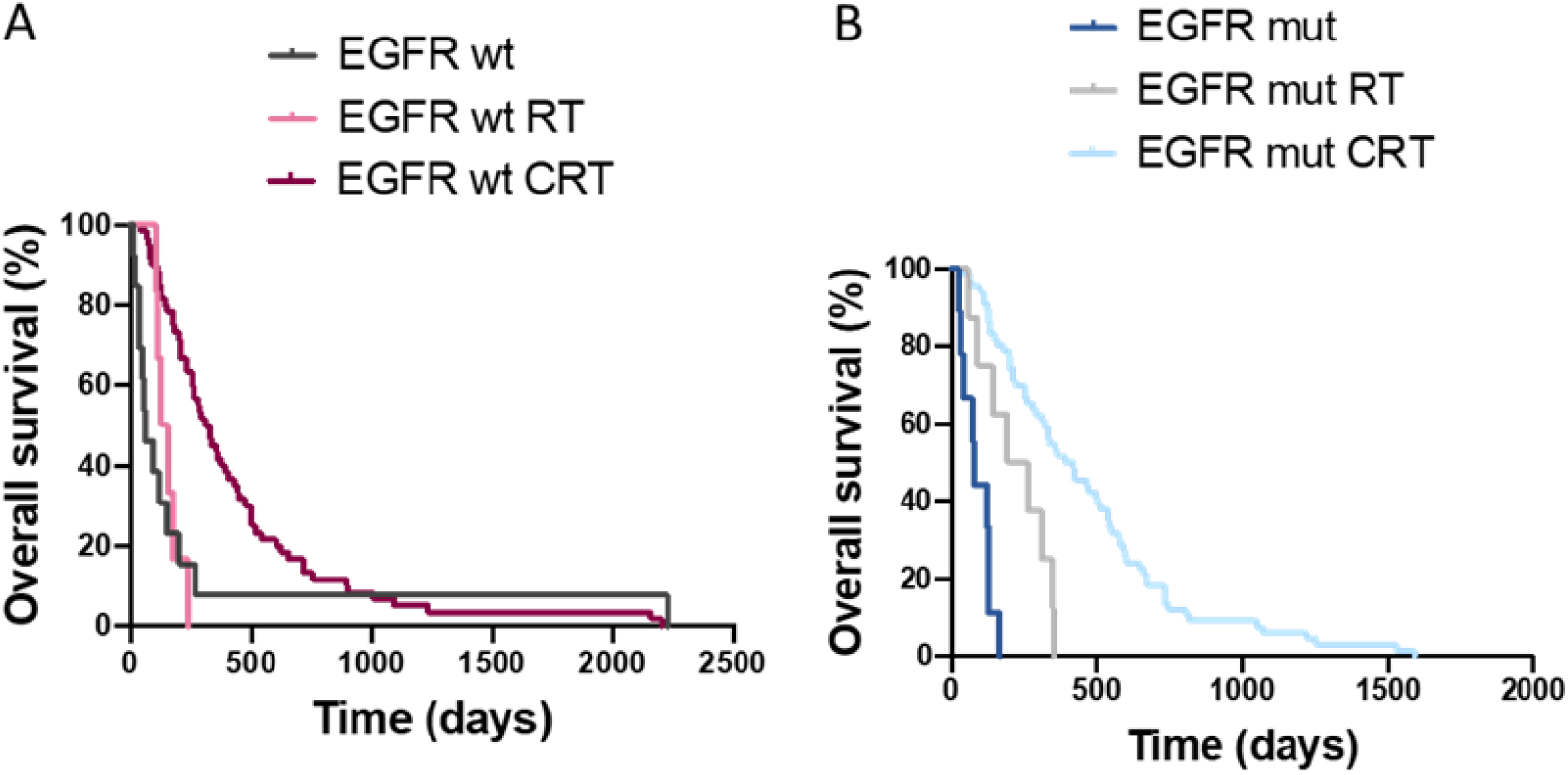
Retrospective analysis in a human data set. Retrospective analysis of survival stratified by EGFR status (EGFRwt (left) and EGFRmut (right)) in human data (TOG database). Kaplan–Meier curves show limited benefit of CRT with respect to RT alone in patients with EGFRwt and a clearer effect in those with EGFRmut.

Among EGFRwt patients treated with CRT, survival after PR was very similar to biopsy (median OS 6.6 vs. 6.4 months; p = 0.45; Suppl. Fig. S2A), whereas only GTR was associated with a modest, non-significant survival gain with respect to PR (median OS 10.7 months; p = 0.21; Suppl. Fig. S2A). By contrast, in EGFRmut patients treated with CRT, increasing surgical extent was consistently associated with progressively better outcomes (median OS 4.3, 11.6, and 19.1 months for biopsy, PR, and TR, respectively; p = 1.03e−05 for biopsy vs. PR; p = 0.23 for PR vs. GTR; Suppl. Fig. S2B).

To further validate our findings, we analyzed the TCGA GBM cohort (TCGA-GBM project, accessed July 1, 2025). As in the TOG dataset, the number of patients treated with TMZ monotherapy after applying the filter of EGFR status was very small (N = 3), precluding statistically meaningful conclusions for this subgroup. Similarly, only a few patients (N = 22, with OS data available for 7) had not received RT, which prevented comparisons between RT and surgery, as performed in the TOG cohort. Nevertheless, EGFR status emerged as a prognostic factor in the full TCGA dataset: consistent with the in vivo results, patients with EGFRwt tumors (N = 162) showed longer OS compared with those harboring EGFRmut tumors. Within the EGFRwt subgroup, patients treated with RT alone (N = 51) had limited survival (median OS 11 months) compared with those receiving CRT with TMZ as chemotherapy (N = 89; median OS 21 months; p = 1.29e−6; Fig. 5c). In EGFRmut tumors, OS was markedly prolonged when TMZ was added to RT (N = 105 for RT only and 300 for CRT; median OS 5 vs. 16 months; p = 1.11e−16; Fig. 5d).

## Discussion

The standard treatment of GBM patients, based on a combination of RT and chemotherapy with TMZ, was established more than 20 years ago. However, it is not curative and can only control the disease for a limited time. In order to improve the Stupp protocol, intensification of TMZ [10–12] or RT [13] delivery schemes has been attempted, without positive results on OS and with increased toxicity observed in certain cases [14, 15]. As an alternative, we have showed recently that longer time spacing between TMZ treatments could be beneficial for slow-growing GBM. Moreover, these spaced chemotherapy reduced the appearance of resistance markers and allowed increased TMZ doses without the associated toxicity [7]. Here, we aimed to determine if enlarging the time interval between RT fractions could also improve the anti-tumor effect in GBM, as we have predicted for lower grade gliomas [16, 17]. For that, we have used the same GBM models used for the TMZ study, though we have not observed a beneficial effect of protracted RT schemes, neither in the anti-tumoral effect nor in the expression of resistance markers in response to irradiation. These results argue in favor of different mechanisms of action for RT and TMZ in GBM and that different models should be used for in silico studies in both cases.

Even though the vIII isoform of EGFR have been linked to radioresistance in gliomas [18, 19], the results presented here indicate that RT can effectively inhibit the growth of the SVZ-EGFRvIII model, but it does not affect the survival of mice carrying SVZ-EGFRwt tumors. Previous studies have evaluated the relationship between the presence of EGFR mutations and the sensitivity to irradiation in lung cancer patients. In agreement with our results in GBM, patients with mutant EGFR respond better to RT, either with locally advanced tumors [20] or with brain metastasis [21]. We hypothesize that differences in the tumor microenvironment of the two models could explain the lack of sensitivity to RT observed in the EGFRwt GBM model. We have previously shown, for example, that in the absence of EGFR mutations, EGFR overexpression in GBM cells leads to the formation of tumors with a disrupted vasculature and more areas of hypoxia [5], which has been linked to radiotherapy resistance in gliomas [22]. Notably, the retrospective analysis of the TOG cohort confirmed that, in patients carrying EGFRwt GBM, the clinical benefit of adding RT to surgery is not significant, whereas in prolongs OS in patients carrying EGFRmut tumors.

Increasing evidence supports the notion that each component of the GBM SOC alters the remaining disease course, including effects on tumor growth characteristics and the tumor microenvironment, emphasizing the difference between treatment-naive and recurrent tumors. However, from the studies compiled in this review, it is also evident that surgical techniques, RT dosing regimens, and TMZ doses and routes of administration are not standardized, making it difficult to compare studies or determine which strategies are optimal for clinical translation

One limitation of our study is that we could not analyze the effect of TMZ alone in the retrospective studies, given the low number of patients treated only with chemotherapy in both the TOG and the TCGA cohort. However, our results suggest that in cases where only one treatment is the clinical option, elucidating the EGFR status could serve to discriminate those that could benefit from RT from those that would rather be treated with TMZ. This is the case for elderly patients, who represent an increasingly common population among GBM patients. Apart from having a worse prognosis [23] , they are less able to tolerate the intensiveness of standard CRT. Hypofractionated radiation plus TMZ or choosing between RT alone over chemotherapy alone are available options for elderly GBM patients [24]. When hypofractionated CRT is intolerable, the recommendation is that patients with unmethylated MGMT promoter should receive RT [25]. Indeed, the phase III NOA-08 trial suggested that the event-free survival was longer in patients with MGMT promoter methylation who received TMZ than in those who underwent RT, whereas the opposite was true for patients with no methylation of the MGMT promoter [26]. Notably, the benefit of TMZ over RT was observed particularly for patients of the RTKII methylation subclass, which is associated with EGFR amplifications, although the presence of additional mutations in the receptor was not analyzed [27]. Our results encourage a retrospective analysis of the data from that trial or the promotion of a new trial for those elderly GBM patients, for whom one of the schedules must be eliminated from the plan, which should be then stratified based on the status of the MGMT promoter and the EGFR gene.

### Authorship

M.C.P., M.P.P., M. I., J.B.B, V.P.G and P.S.G. conceived and planned the experiments; V.P.G and P.S.G got funding; M.C.P., M.P.P., M. I., I.G.S., and A.C. carried out the experiments; M.C.P., M.P.P., M. I., J.B.B, V.P.G and P.S.G analyzed the data and contributed to the interpretation of the results; M.C.P. and P.S.G. took the lead in writing the manuscript. All authors provided critical feedback and contributed to the final document.

### Data availability

The data that support the findings of this study and the materials used are available on request from the corresponding authors.

## Acknowledgements

We sincerely thank the patients and their families to their support of this project.

