## Supplementary Figures for "EGFR-Driven Phenotypes Dictate Differential Therapeutic Response to Radiotherapy and Temozolomide in Glioblastoma"

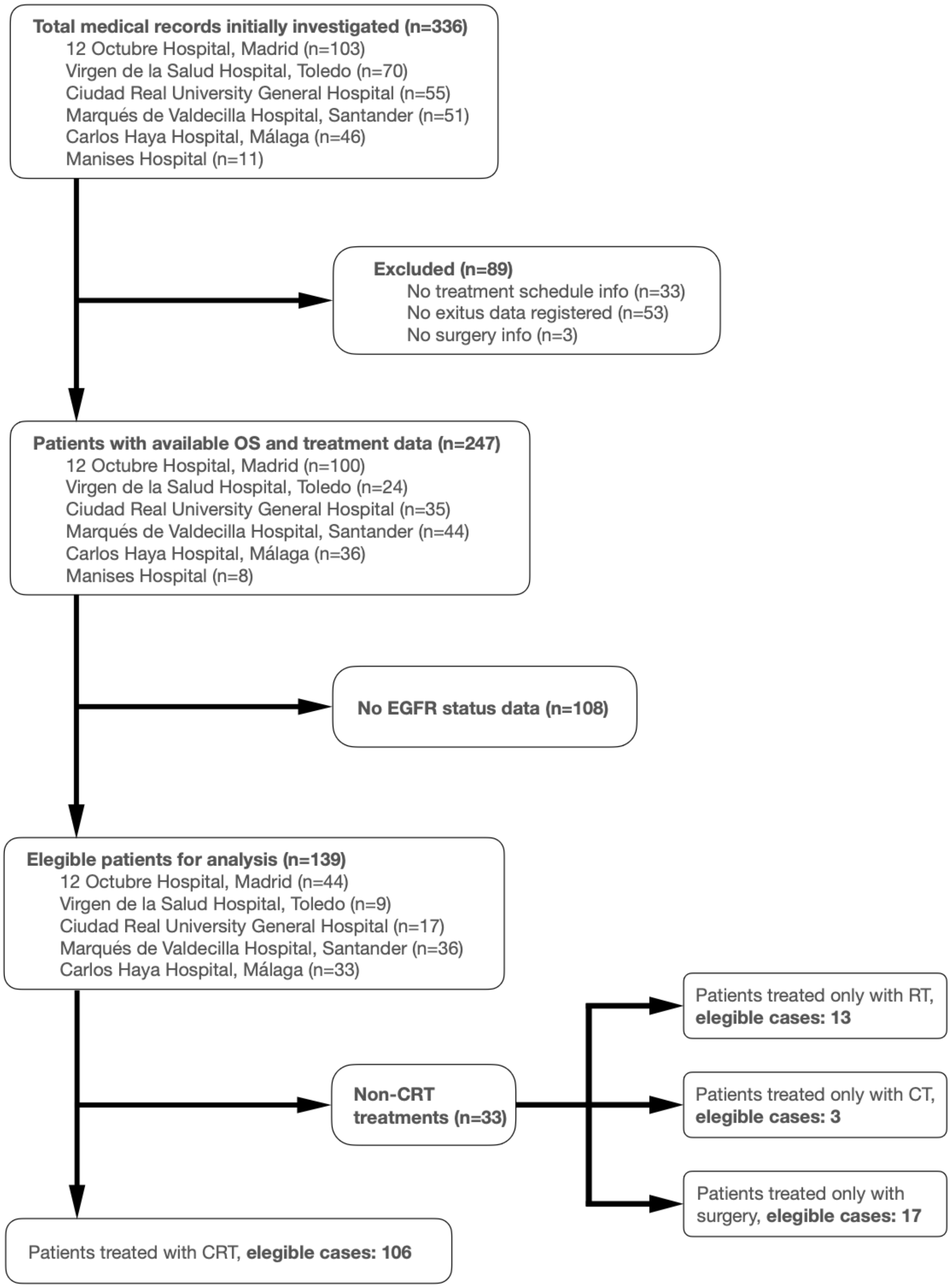


**Supplementary Figure 1. Patient selection and molecular profiling workflow.** Summary of case filtering across participating centers and inclusion criteria. Molecular characterization (Ki67, MGMT promoter methylation, IDH and EGFR status by IHC) was performed following standard protocols. Patients with available EGFR data (n=139) were stratified by treatment group (CRT, RT only, or surgery only).

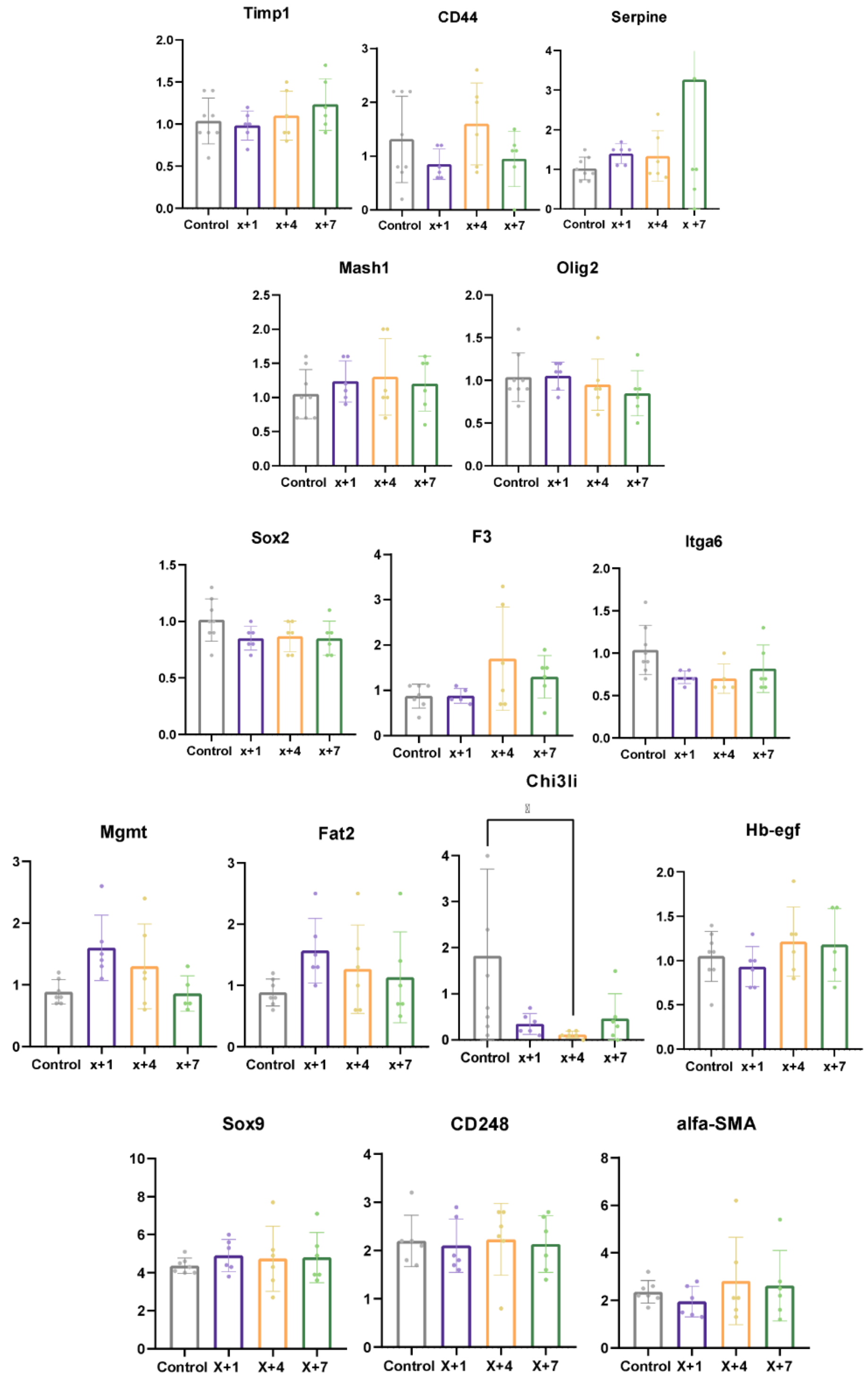


**Supplementary Figure 2.** Transcriptional analysis of phenotypic markers following RT treatment. qRT-PCR analysis of genes associated with mesenchymal, proliferative, persistence, RT-resistance, pericyte, and hypoxia-related phenotypes in SVZ-EGFRvIII tumors collected after the different radiotherapy schedules (Control, X+1, X+4, X+7). Actin expression was used for normalization. *$p < 0.05$, **$p < 0.01$, ***$p < 0.001$.

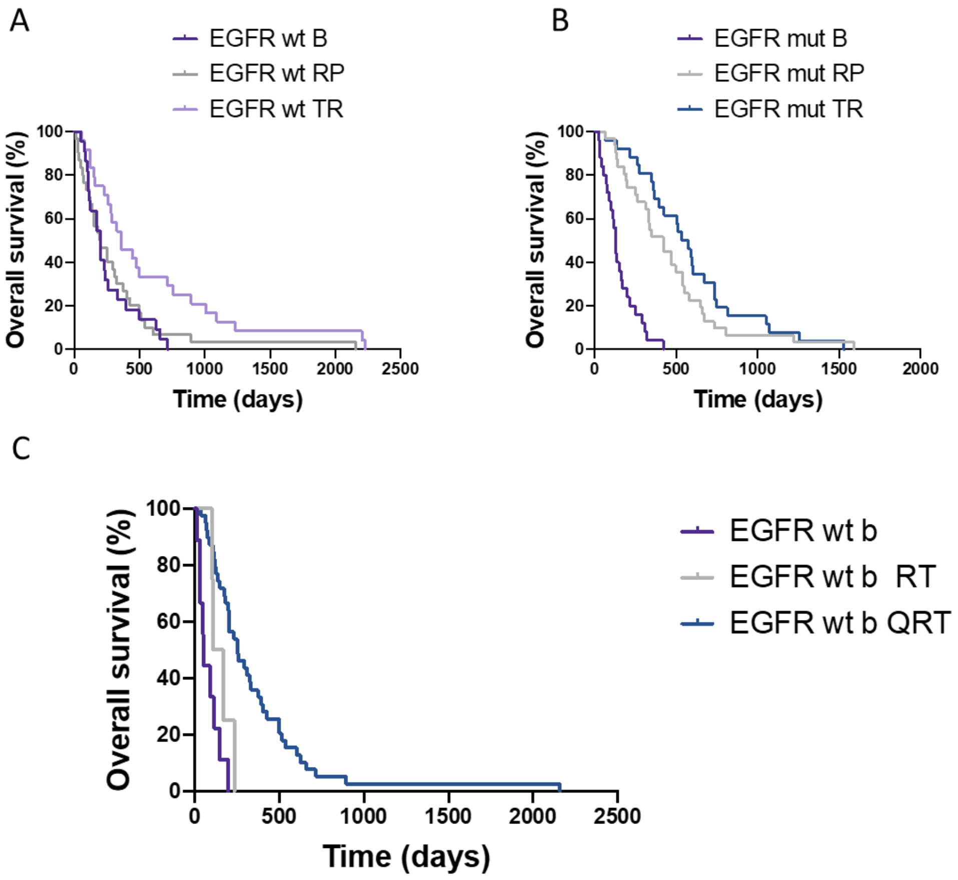


**Supplementary Figure 3. Survival analyses in the TCGA cohort stratified by EGFR status, surgical extent, and treatment**. Kaplan–Meier curves show overall survival in EGFRwt patients treated with CRT (chemo and radiotherapy) according to surgical extent **(A)**, EGFRmut patients treated with CRT **(B)**, and EGFRwt patients undergoing biopsy or PR (partial resection) and subsequently treated with surgery only, RT alone, or CRT **(C)**. Median OS and pairwise p-values are reported in the main text.
